# Targeting influenza at the topologically conserved substructures

**DOI:** 10.1101/2020.10.25.351643

**Authors:** Zubair Ahamed, Kamjula Vandana, Kakunuri Bhuvaneswari

## Abstract

H9N2 avian influenza virus is a low pathogenic endemic strain in the domestic poultry of most of the Asian countries. Attempts have extensively failed in eradicating its diverse strains. To find the drug against the evolutionarily conserved substructures, the target protein sequence is analyzed through sequence and modelled structure for mapping the structurally conserved topology. The available drugs are screened against the deciphered topological map through the predicted ADMET and drug-likelihood scores. This study helps to build a theoretical framework to make the foremost potent drug.

## 1. Introduction

Influenza virus belongs to the Orthomyxoviridae family and is known to cause respiratory tract infection (Flu) in vertebrates (Mourya, et al., 2019). It is categorized into four subtypes, namely A, B, C and D on the basis of its antigenic differences among nucleoprotein (NP) and matrix (M) protein (Nicole M. and Peter, 2008). Although the influenza A virus spreads naturally, a few of its viral strains have the capability to replicate in the host organisms, like horses and pigs, and are known to subsequently infect humans (Taisuke and Kawaoka, 2001). While the influenza B virus is less pathogenic and is localized to humans (Bodewes, et al., 2013), the influenza C infects swine, dogs and humans and is a rare virus category (Bailey, et al., 2018). However, the influenza D strain is only found to infect cattle (Su, et al., 2017).

Influenza virus is an RNA virus that is subject to several point mutations across its antigenicity-determining region. As the virus evades pro-existing immunity of a host, it leads to annual/occasional pandemic conditions. For a higher rate of point mutations, genetic segments often get reassorted to form an unprecedented viral strain (Lu, et al., 2018). Hence, the virus has been posing a significant threat to mankind for several years.

H9N2 is a subtype of influenza A virus which is pandemic in poultry and acts as a gene donor for the subtypes viz., H7N9 and H10N8 virus which affects mammals (Chengjun, et al., 2005). It is prevalent across many countries in Asia, Africa, Europe and Middle East (Pusch and Suarez, 2018). On the basis of its pathogenicity, the influenza virus has a unique capacity to produce different subtypes by antigenic drift/shift. It is usually categorized into two types, viz. high pathogenic avian influenza virus (HPAIV) (Swayne, 2007) with a mortality rate above 95% and low pathogenic avian influenza virus (LPAIV) (Kwon, et al., 2008) that leads to respiratory failure, pneumonia, bronchitis and asthma attack. On the basis of several *in-vivo* studies, the pathogenicity of H9N2 is found to be low (Rui-Hua, et al., 2011). While H9 stands for hemagglutinin (HA) and is a glycoprotein responsible for the internalization of the virus by binding to the host receptor (Stevens, 2004), N2 defines the neuraminidase subtype, bearing the multifunctional neuraminidase/sialidase on the virus envelope. The HA protein is a homotrimer and has a receptor pocket over the globular head of every chain. For viral activation, it must be cleaved by trypsin-like serine endoprotease between the HA1 and HA2 domains (Constance, et al., 1986). Further, the neuraminidase protein is required in all stages of viral infection and is primarily significant for the release of virions from the infected cells (Shtyrya, et al., 2009). Till date around 18 HA (H1-H18) subtypes have been identified, and H17 and H18, H17N10 and H18N11 have recently been found in bats, fruit bats and peruvian bats respectively (Mehle, 2014). Further, 11 NA(N1-N11) subtypes have been identified, and it results in hundreds of diverse viral combinations (Gao, et al., 2010). Firstly found in 1874 in Italy as a causative agent for a massive poultry death, H9N2 is recently shown to evolutionarily diverge into several other influenzae subtypes namely, H5N1, H7N9, H10N8 and H5N6 (Fusaro, et al., 2011; Jian-Hong, et al., 2005).

For efficiently targeting all influenza species through a single drug, the diverse, it is important to analyze the H9N2 sequence and map the evolutionarily conserved segments across the key site(s). It is because the key functional site is said to encode a high degree of conservation for most of the key residues, defining its spatial topology, and the drugs blocking such crucial residues should be the most potent drug. The structural and functional study of H9N2 virus is thus found to be crucial to identify its role in the reassortment of the genes in the lethal subtypes (Pu, et al., 2015). The consensus based prophylactic approaches have already been investigated to design a novel antigen from broadly reactive antigen (Carter, et al., 2016; Wong, et al., 2017) against influenza. However, a detailed computational analysis has never been implemented for the HA drugs so that on basis of their interaction fingerprint and the key binding residues, a more potent drug molecule could be designed. The study would pave the way for the development of an efficient drug molecule against the conserved substructure of the diverse virus strains.

## 2. Materials and Methods

### 2.1 Sequence analysis

The 560-residue HA protein sequence of influenza A virus (H9N2), with accession number AWK91180.1 is retrieved from NCBI protein database. For domain, motif and confirmation of the functional details, NCBI CDD, Pfam and TIGRFAM15.0 databases are orderly screened through the HHPred tool (Zimmermann, et al., 2018), and the functionally conserved signature sequences are subsequently scanned within the protein databases through InterProScan5 (Madeira, et al., 2019). The important physicochemical properties, viz. atomic and residue composition, molecular weight, estimated half-life, extinction coefficient, aliphatic index, theoretical isoelectric point (pI), and instability index are lastly estimated for the selected sequence through ProtParam (Gasteiger, et al., 2005).

### 1.2. Constructing the reliable model conformation

As the HA protein structure is not available in the protein data bank (PDB), it is predicted through the template based modeling strategy (Ashish and Shibasish, 2013; Runthala, 2012; Runthala and Chowdhury, 2016; Runthala and Chowdhury, 2019). Its hidden markov model (HMM) based sequence profile is constructed through HHPred (Zimmermann, et al., 2018) against the PDB database, and the top-ranked functionally similar templates are assessed through the sequence and structural similarity scores (Runthala and Chowdhury, 2019), using the in-house scripts.

The top-ranked template, sharing a functional similarity with the input sequence, is selected for constructing the HA model, as discussed earlier (Runthala, 2020; Runthala and Chowdhury, 2014; Shikhin, et al., 2016; Sunil and Ashish, 2016). The alignment is curated and the structure is predicted through MODELLER9.22. To relieve the atomic clashes and construct the best possible near-native model (Shikhin, et al., 2016), the predicted structure is energetically relaxed through iterative sampling rounds through 3drefine. The sampled model is topologically assessed through TM-Score (Zhang and Skolnick, 2004) and root mean squared deviation (RMSD) against the deployed template.

### 1.3 Mapping the structural conservation

To extract the structurally conserved substructures and decipher the most crucial segments in the HA protein for further targeting them through the best drugs, consurf algorithm is utilized (Ashkenazy, et al., 2016). To strategically approach the issue a better way, HMMER is used to iteratively search the UNIREF90 database through five iterative rounds for constructing the sequence profile of HA protein. To extend the search to the maximum set of functionally similar sequences, the alignment is constructed by HMMER through iterative rounds with an E-value threshold of 0.0001. The algorithm is designed to maximally sample the homologues within a mutual sequence identity of 35-95%. MAFFT-L-INS-i algorithm is used to align the screened homologues and using the Bayesian analysis, the best model is considered as the evolutionary substitution model. It is expected to robustly compute the conserASY server (www.expasy.ch/tools) (Artimo, et al., 2012).

### 1.4 Screening the top-ranked drug molecules

As per the recent research articles, the potent drug candidates, shown to be active against the HA protein of virus, are selected. This dataset includes viz. N-Cyclohexyltaurine, Arbidol, Rimantadine, Amantadine, Oleanane acid, Dimethylbenzanthracene and Baloxavir marboxil, as shown with the two-dimensional maps in Figure1. As the protein-ligand docking has been traditionally deployed to computationally screen the potent drug molecules, the screened dataset is docked through the AutoDockTools (Morris, et al., 2009). The receptor conformation of the predicted HA model is prepared by adding the hydrogen atoms and saving it in the .pdbqt format. All the ligand molecules are subsequently prepared in the required manner and saved in the .pdbqt format as well. Lamarckian genetic algorithm, an adaptive local search protocol, is subsequently deployed to guide the docking simulation with a maximum number of 2.5 million energy evaluations through utmost 27,000 generations. To reliably guide the conformational scan within the ligand-topology, the top-ranked cavity of the predicted model is localized through Prankweb protocol (Jendele, et al., 2019). The model-atom I276-CA is selected as the gridcenter with a 50×50×50 gridbox to optimally cover the interaction zone. The lowest energy docking solution is considered to be the near-native protein-ligand conformation.

**Figure1:**
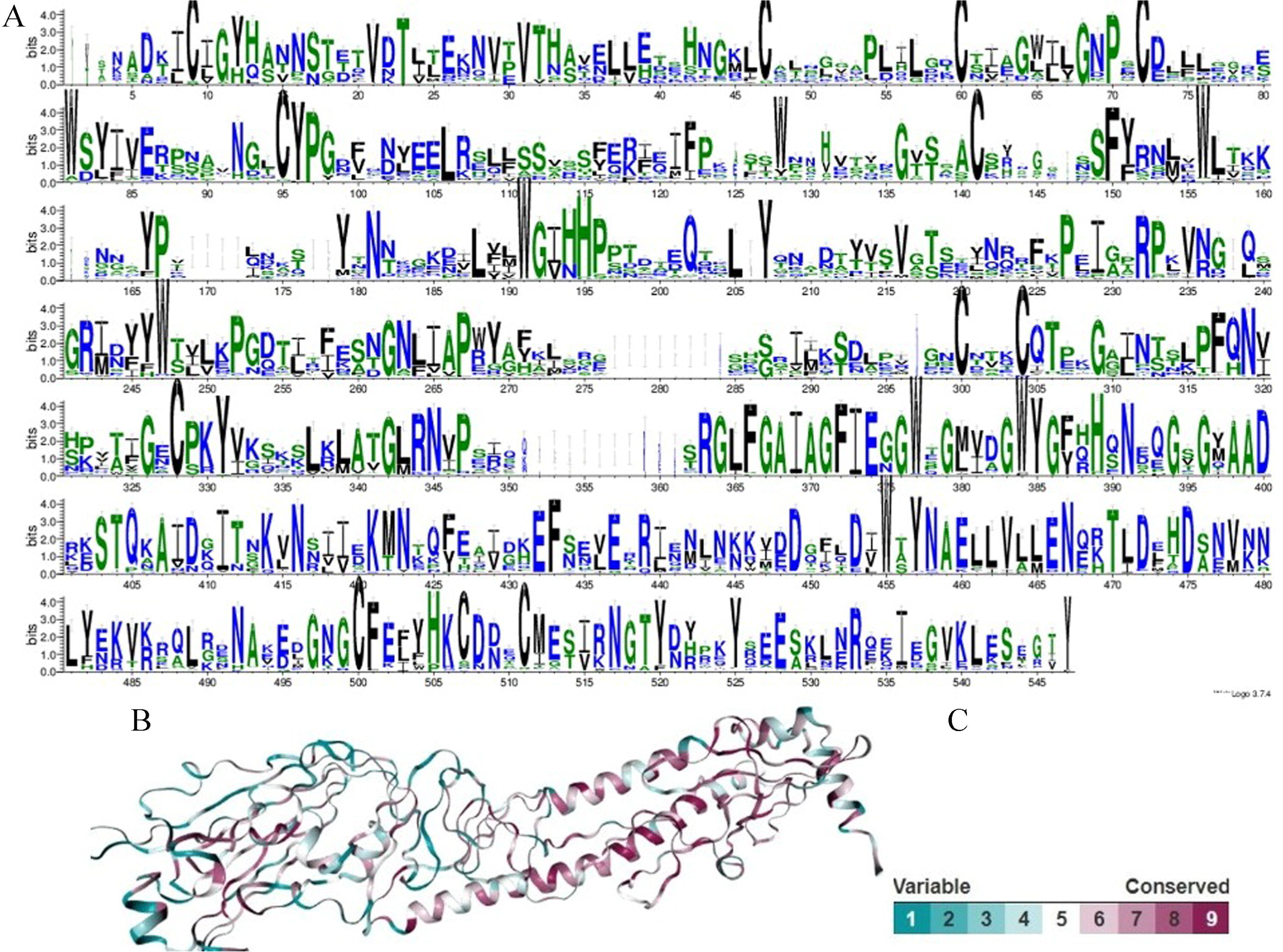
Two-dimensional representations of the seven retrieved drugs.

### 1.5 ADMET profile and drug-like prediction

Many potential therapeutic agents fail to reach the clinic trials due to their biologically unfavorable absorption, distribution, metabolism, elimination and toxicity (ADMET) parameters and drug-likeness score (Siramshetty, et al., 2016), and to find the efficacious drugs and to minimize the undesired effects, it is important to evaluate such properties. The SwissADME (Daina, et al., 2017) and Molinspiration (Ertl, et al., 2000) algorithms are used to predict these properties for the drug molecules. To select the best drug molecule, several pharmacokinetic properties, viz. miLogP, topological polar surface area (TPSA), molecular weight (MW, g/mol), hydrogen bond donors (nON), hydrogen bond acceptors (nONH), number of rotatable bonds (nrotb), volume, uptake in gastrointestinal cells (GI absorption) and water solubility are estimated. Moreover, the molecules are evaluated through the Lipinski’s thumb rules (Lipinski, 2004) to select the most potent drug molecule.

## 2. Results and Discussion

### 2.1 Sequence analysis

To estimate the key physico-chemical parameters for the selected sequence AWK91180.1, Protparam is used (Gasteiger, et al., 2005). The server allows the theoretical estimation of several parameters, viz. molecular weight, isoelectric point, residue composition, extinction coefficient, estimated half-life, instability index and grand average of hydropathicity (GRAVY).

The resultant scores indicate that molecular weight of HA protein is 62.641KDa, and its theoretical isoelectric point is 7.18, which further shows that the protein should be neutral in nature. Within the 560-residue selected protein, the negatively and positively charged residues are orderly found to be 56 and 56, and it indicates that the net charge of the protein is neutral, as also expected. The extinction coefficient for the selected protein is found to be 90145, and it represents the amount of light absorbed by the protein at a certain wavelength. The value is experimentally utilized to determine the concentration of protein in solution for pursuing its purification. To estimate the stability of the protein in a test tube, its instability index is estimated. It is found to be 32.21, and as the score is lower than 40, the protein is found to be highly stable.

By screening the conserved domain database (CDD), protein family and TIGR family database (TIGRFAM15.0) with the deployment of hidden markov model (HMM) based sequence profiles, and by searching the InterProScan5 server for the considered sequence, functional details of the considered sequence is affirmed. It is interesting to observe that the 14-522 sequence chunk is topologically as well as functionally conserved across the sequence and structural databases.

### 2.2 Constructing the reliable model conformation

Through HHPred, the 11 functionally similar templates are found to maximally cover the 560-residue target sequence. Through the *in-house* scripts (Runthala and Chowdhury, 2019), the structures are ranked against the target sequence. The top-ranked 509-residue template 4F23A, bearing an E-value of 3.0E^−209^, shows a sequence identity of 51.41% and is used for modelling the selected HA protein. However, homology for the N- and C- terminal sequence is found to be substantially lower against the functionally similar structures, and these terminal segments (1-18 and 517-560) are removed from the input sequence, as usually done by several modelling servers including Swissmodel (Biasini, et al., 2014).

On basis of the selected template 4F23A, the curated target-template alignment is structurally assessed by Espript3.0 (Figure 2A) (Robert and Gouet, 2014) and it shows a complete overlap for the secondary structure elements. The alignment is subsequently used for building a near-native structure through MODELLER9.22 (Webb and Sali, 2016). Iterative sampling methodology has been shown to be more accurate than the usual sampling methodology, and is deployed to alleviate the non-physical atomic clashes and to build a more accurate model topology (Runthala and Chowdhury, 2014). Within this 498-residue protein structure, besides the flanking unaligned C-terminal segment, the conserved core of 497 residues is found to be topologically overlapped within a distance deviation of 5Å, as shown by TM-align (Figure 2B). The model orderly shows a TM-score and RMSD score of 0.997 and 0.19 against the template, and with an ERRAT score (Chris and Todd O, 1993) of 92.7536, it is validated to be a near-native structure. The model further shows a QMEAN score of −0.44, and it defines the likelihood of the local model topology to its native structure (Figure 2C). Evaluating the Ramachandran map of the predicted model, it is found that 94.49% residues are found to be placed within the favored regions (Figure 2D). Moreover, the Molprobity score of 1.64 is found for the predicted structure, and its topology is confirmed to be a credible one. The predicted model is subsequently docked with the seven reported ligands (Figure 2E).

**Figure 2.**
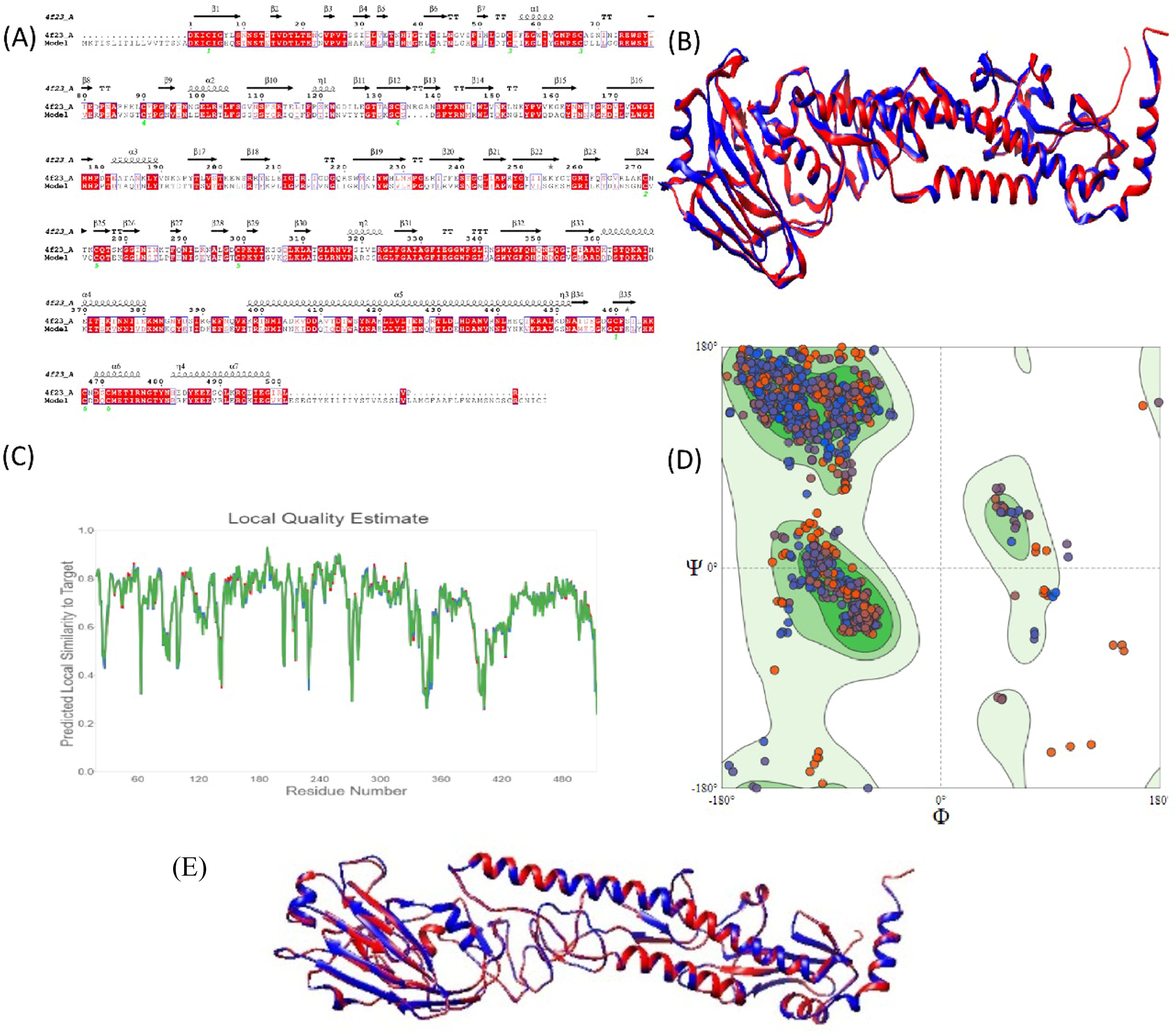
(A) Espript3 result for the curated sequence alignment decodes the conserved residues in red sequence block, secondary structure elements as squiggled helices, arrow-marked beta strands and the TT-labeled turns (B) Structural overlap of the HA protein (AWK91180.1) from influenza A virus (Blue) against the 4F23A structure (Red), and highlighting the high-scoring conserved topology (C) QMean assessment (D) Ramachandran map of the predicted model. (E) Structural superimposition of the predicted model and template.

### 2.3 Mapping the structural conservation

ConSurf algorithm is deployed to determine the conservation at every sequence position for the considered sequence. It statistically computes the conservation score for each residue on basis of the multiple sequence alignment for the screened set of sequence homologues. It is widely used to detect the functional loci on proteins as the residues, often found conserved across the phylogenetic lineage.

UNIREF90 database is screened through 3 rounds of PSI-BLAST at an E-value threshold of 0.0001, and excluding the functional outliers, the top-ranked 150 closest orthologues (*including the input sequence*), sharing a sequence identity of 35-95% (Supplementary table 1). Sequence profile is constructed through hidden markov model based Clustal omega alignment, and the conservation profile for the evolutionarily retained sites is drawn for the conserved HA model (Zimmermann, et al., 2018). Consensus sequence is computed from this alignment to analyze the degree of conservation for the evolutionarily retained sites. MSA of the theoretically constructed, conserved HA structure and the top-ranked homologues is used to estimate the evolutionary conservation score across the chain through Weblogo (Crooks, et al., 2004), as represented in Figure3A. It shows a significantly high conservation of residues across the chain. To structurally localize the conservation, empirical Bayesian method is deployed to estimate the rate of evolution and it is further used to measure the residue conservation across the chain through Consurf. The conservation pattern is plotted on the constructed protein model, and is then visualized by using Chimera.

**Figure 3:**
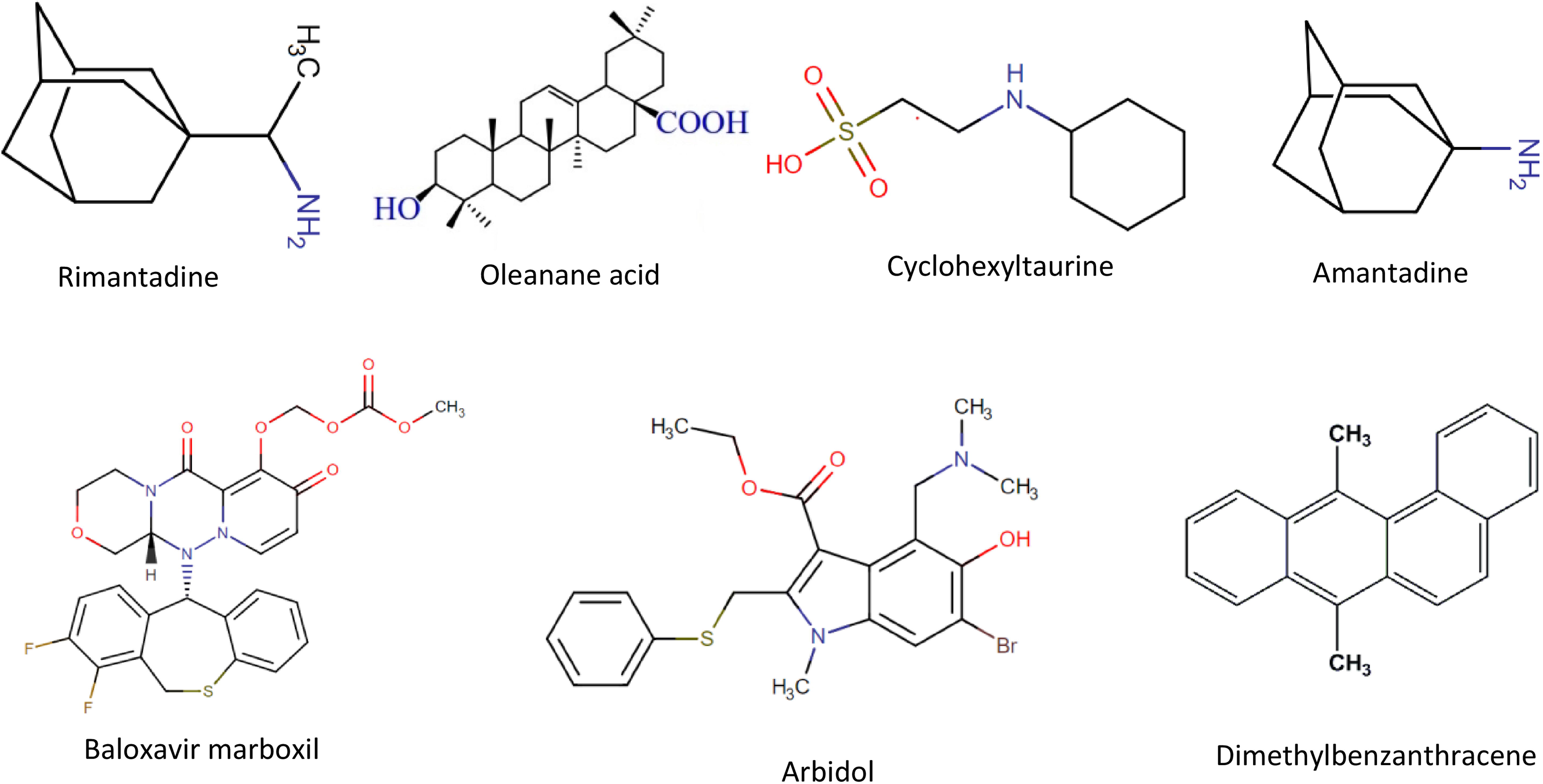
(A) Conservation logo of AWK91180.1 for the sequence dataset, indicating a complete preservation for several loci (B) Evolutionary conservation of the considered HA protein (AWK91180.1) structure (C) Standard color-grading deployed for the analysis.

Sequence conservation for the highly conserved, average conserved and variable positions is orderly represented through maroon, white and turquoise colors on the protein surface (Figure3B), as per the standard color-grading (Figure3C). The analysis shows an average pairwise distance score of 0.88586. It shows a substantially low range of sequence conservation from 0.0213 to 1.859. Further, 44 positions, bearing the highest grade point of 9, are found to be completely conserved within the constructed profile. Moreover, it also shows that out of a total of 560 residues, 74, 76 and 73 residues score a consurf grading of 8, 7 and 6 respectively, and it thus shows that 47.67% residues within the entire HA structure are greatly conserved and should be functionally important.

### 2.4 Screening the top-ranked drug molecules

To find the top-ranked active site of the constructed protein model for the selected HA protein (AWK91180.1) from influenza A, the prankweb server is used and accordingly the interaction zone is deciphered. The I276-CA atom is subsequently selected as the gridcenter within this region and the ligand-dataset is docked in the top-ranked site. The top-ranked antagonist is selected on the basis of their inhibition constants (Ki) and interaction energy (ΔG) against the model. Using Autodock, the molecular interaction energy and Ki scores of the ligands are estimate, and Baloxavir marboxil is found to be the most potent lead compound (Table1), and the results are in line with the recently published research (Fukao, et al., 2019). The drug shows a significantly low energetic score of −9.32Kcal/mol, along with a very low Ki score of 146.33nM. The interaction plot is subsequently constructed for the docked complexes of these ligands through Ligplot+ (Laskowski and Swindells, 2011), as shown in the following Figure4. Further, all the shortest distance between the interacting residues and the ligand atom is calculated through Chimera, as enlisted in Table1.

**Table1:** The binding energy, Ki score and the closest intermolecular bond length between the atoms of the known drug molecules and the predicted HA structure.

| # | Drug | Binding energy (Kcal/mol) | Ki | Interacting residues along with their distances |  |
| --- | --- | --- | --- | --- | --- |
| 1 | Baloxavir marboxil | -9.32 | 146.33nM | 1 | THR74-O7 : UNK 900-C26 [1.223 Å°] |
|  |  |  |  | 2 | GLU76-O6 : UNK 900-C26 [1.363 Å°] |
|  |  |  |  | 3 | GLU99-O7 : UNK 900-C26 [1.223 Å°] |
|  |  |  |  | 4 | ARG100-C18 : UNK 900-O4 [1.226 Å°] |
|  |  |  |  | 5 | PRO101-C17 : UNK 900-C11 [1.336 Å°] |
|  |  |  |  | 6 | ARG120-C5 : UNK 900-C6 [1.506 Å°] |
|  |  |  |  | 7 | ARG275-C24 : UNK 900-C20 [1.393 Å°] |
|  |  |  |  | 8 | ILE276-C2 : UNK 900-N1 [1.466 Å°] |
|  |  |  |  | 9 | LEU277-C11 : UNK 900-N3 [1.383 Å°] |
|  |  |  |  | 10 | LYS278-C16 : UNK 900-C22 [1.392 Å°] |
|  |  |  |  | 11 | TYR309-C22 : UNK 900-C24 [1.383 Å°] |
|  |  |  |  | 12 | ASP405-C12 : UNK 900-S1 [1.796 Å°] |
|  |  |  |  | 13 | HIS406-C23 : UNK 900-F2 [1.339 Å°] |
|  |  |  |  | 14 | GLU407-N1 : UNK 900-C1 [1.462 Å°] |
|  |  |  |  | 15 | PHE408-C4 : UNK 900-N2 [1.458 Å°] |
|  |  |  |  | 16 | ASN416-C19 : UNK 900-F1 [1.341 Å°] |
| 2 | Oleanane Acid | -7.96 | 1.47 µM | 1 | THR74-C20 : UNK 900-C17 [1.527 Å°] |
|  |  |  |  | 2 | GLU76-C17 : UNK 900-O1 [1.427 Å°] |
|  |  |  |  | 3 | GLU99-C13 : UNK 900-C20 [1.527 Å°] |
|  |  |  |  | 4 | PRO101-C18 : UNK 900-C7 [1.349 Å°] |
|  |  |  |  | 5 | ARG120-H47 : UNK 900-O1 [0.972 Å°] |
|  |  |  |  | 6 | ILE276-C18 : UNK 900-C11 [1.498 Å°] |
|  |  |  |  | 7 | ARG275-C30 : UNK 900-C24 [1.540 Å°] |
|  |  |  |  | 8 | LYS278-C10 : UNK 900-C7 [1.532 Å°] |
|  |  |  |  | 9 | GLU407-C8 : UNK 900-C6 [1.532 Å°] |
|  |  |  |  | 10 | PHE408-C8 : UNK 900-C6 [1.532 Å°] |
| 3 | Arbidol | -7.55 | 2.93 µM | 1 | GLU99-C2 : UNK 900-N1 [1.377 Å°] |
|  |  |  |  | 2 | ARG120-C22 : UNK 900-C21 [1.396 Å°] |
|  |  |  |  | 3 | ARG275-C11 : UNK 900-BR1 [1.899 Å°] |
|  |  |  |  | 4 | ILE276-S1 : UNK 900-C9 [1.824 Å°] |
|  |  |  |  | 5 | LYS278-C10 : UNK 900-N1 [1.449 Å°] |
|  |  |  |  | 6 | TYR309-C11 : UNK 900-C8 [1.395 Å°] |
|  |  |  |  | 7 | ASP405-N2 : UNK 900-C14 [1.463 Å°] |
|  |  |  |  | 8 | HIS406-C12 : UNK 900-O2 [1.380 Å°] |
|  |  |  |  | 9 | GLU407-C3 : UNK 900-C4 [1.376 Å°] |
|  |  |  |  | 10 | PHE408-C19 : UNK 900-C16 [1.514 Å°] |
|  |  |  |  | 11 | ASN416-N2 : UNK 900-C13 [1.463 Å°] |
| 4 | Rimantadine | -7.43 | 3.57 µM | 1 | ARG275-N1 : UNK 900-H21 [1.019 Å°] |
|  |  |  |  | 2 | ILE276-N1 : UNK900-C11 [1.461 Å°] |
|  |  |  |  | 3 | LEU277-C9 : UNK 900-C4 [1.538 Å°] |
|  |  |  |  | 4 | LYS278-C10 : UNK 900-C2 [1.538 Å°] |
|  |  |  |  | 5 | TYR309-C10 : UNK 900-C3 [1.538 Å°] |
|  |  |  |  | 6 | ASP405-C1 : UNK 900-C11 [1.531 Å°] |
|  |  |  |  | 7 | HIS406-C12 : UNK 900-C11 [1.525 Å°] |
|  |  |  |  | 8 | GLU407-N1 : UNK 900-C11 [1.461 Å°] |
| 5 | dimethylbenzanthracene | -6.83 | 9.88 µM | 1 | GLU99-C18 : UNK 900-C14 [1.395 Å°] |
|  |  |  |  | 2 | ARG120-C10 : UNK 900-C9 [1.391 Å°] |
|  |  |  |  | 3 | ILE276-C17 : UNK 900-C18 [1.391 Å°] |
|  |  |  |  | 4 | LEU277-C17 : UNK 900-C13 [1.395 Å°] |
|  |  |  |  | 5 | LYS278-C19 : UNK 900-C15 [1.397 Å°] |
|  |  |  |  | 6 | TYR309-C19 : UNK 900-C15 [1.397 Å°] |
|  |  |  |  | 7 | ASP405-C19 : UNK 900-C20 [1.387 Å°] |
|  |  |  |  | 8 | HIS406-C16 : UNK 900-C8 [1.405 Å°] |
|  |  |  |  | 9 | GLU407-C18 : UNK 900-C14 [1.395 Å°] |
|  |  |  |  | 10 | PHE408-C10: UNK 900-C9 [1.391 Å°] |
|  |  |  |  | 11 | ASN416-C10 : UNK 900-C8 [1.395 Å°] |
|  |  |  |  | 12 | ASN420-C20 : UNK 900-C16 [1.392 Å°] |
| 6 | Amantadine | -6.54 | 16.13 µM | 1 | ARG275-C4 : UNK 900-C7 [1.540 Å°] |
|  |  |  |  | 2 | ILE276-C8 : UNK 900-C3 [1.538 Å°] |
|  |  |  |  | 3 | TYR309-C3 : UNK 900-C6 [1.540 Å°] |
|  |  |  |  | 4 | ASP405-N1 : UNK 900-C1 [1.461 Å°] |
|  |  |  |  | 5 | HIS406-N1 : UNK 900-H17 [1.020 Å°] |
|  |  |  |  | 6 | GLU407-C10 : UNK 900-C4 [1.538 Å°] |
| 7 | N-cyclohexyltaurine | -5.02 | 210.17 µM | 1 | ARG275-C4 : UNK 900-C2 [1.529 Å°] |
|  |  |  |  | 2 | ILE276-N1 : UNK 900-H12 [1.021 Å°] |
|  |  |  |  | 3 | LEU277-N1 : UNK 900-C7 [1.456 Å°] |
|  |  |  |  | 4 | LYS278-S1 : UNK 900-O2 [1.451 Å°] |
|  |  |  |  | 5 | GLU293-H17 : UNK 900-O1 [0.987 Å°] |
|  |  |  |  | 6 | TYR309-C8 : UNK 900-C7 [1.520 Å°] |
|  |  |  |  | 7 | ASP405-C6 : UNK 900-C5 [1.526 Å°] |
|  |  |  |  | 8 | HIS406-C6 : UNK 900-C4 [1.526 Å°] |
|  |  |  |  | 9 | GLU407-N1 : UNK 900-C1 [1.460 Å°] |

Figure4 and Table1 uncover a lot of crucial information about the interaction fingerprint of this HA protein. It is found that GLU407 and ILE276 interact with all the drug molecules. Moreover, only ARG100 is found to interact with the most potent Baloxavir marboxil molecule, showing the lowest ΔG and Ki score of −9.32Kcal/mol and 146.33nM, and at a distance of 1.226Å, and is probably the reason that substantially increases the affinity over all the other ligands. Likewise, the residues THR74, GLU76 and PRO101 interact with the two most active drug molecules, viz. Baloxavir marboxil and Oleanane Acid, and probably stabilize the energetic interactions with both each of these ligands.

Figure4: Molecular interaction profile and pharmacophore features of the selected ligands with the active site of the influenza protein.

### 2.5 ADMET profile and drug-like prediction

To select the most efficient drug molecule, SwissADME and Molinspiration algorithms are used to predict the key pharmacokinetic properties for the selected drug molecules (Table2). To affirm the credibility of a potent drug, Lipinski rule is used for the selected drugs, and it states that a drug-like molecule should have logP<5, molecular weight<500 Dalton, and utmost 10 and hydrogen bond acceptors and donors respectively (Lipinski, 2004). As per the allowed range of a potent lead molecule, our compounds show a reasonable water solubility and topological polar surface area (TPSA) (Table 4). The analysis indicates that the top-ranked three drugs, viz. Baloxavir marboxil, oleanane acid and Arbidol with the lowest binding energy and Ki score against the receptor, violates the Lipinski’s rules for only 1 of the required properties. Although these lead-like molecules are not estimated to be highly soluble, their TPSA is found to be on the higher side, and Baloxavir marboxil shows the higher TPSA score of 99.56. It implies that the drug should have a high bioavailability and absorption, as shown earlier (Ertl, et al., 2000).

**Table2:** Scoring parameters for the drug. TPSA: Total Polar Surface Area, MW: Molecular Weight, miLogP: Octanol-water partition coefficient, nON: Number of hydrogen bond donors, nOHNH: Number of hydrogen bond acceptors, nrotb: Number of rotatable bonds

| S. NO. | Drug name | Lipinski rule of 5 | Water solubility | TPSA | MW | nON | nOHNH | nrotb | volume | GI Absorption |
| --- | --- | --- | --- | --- | --- | --- | --- | --- | --- | --- |
| 1 | Arbidol | 0 violation | Moderately soluble | 54.70 | 477.42 | 5 | 1 | 8 | 386.24 | High |
| 2 | Dimethylbenz anthracene | 1 violation | Moderately soluble | 0.00 | 256.35 | 0 | 0 | 0 | 249.14 | Low |
| 3 | Baloxavir marboxil | 1 violation | Poorly soluble | 99.56 | 571.56 | 10 | 0 | 6 | 460.38 | High |
| 4 | Oleanane acid | 1 violation | Poorly soluble | 57.53 | 456.71 | 3 | 2 | 1 | 471.14 | Low |
| 5 | Amantadine | 0 violation | Soluble | 26.02 | 151.25 | 1 | 2 | 0 | 159.20 | High |
| 6 | Rimantadine | 0 violation | Soluble | 26.02 | 179.31 | 1 | 2 | 1 | 192.58 | High |
| 7 | Cyclohexyl taurine | 0 violation | Highly soluble | 66.40 | 207.29 | 4 | 2 | 4 | 188.09 | High |

The three top-ranked drugs have 10, 5 and 3 hydrogen bond donors and 0, 1 and 2 hydrogen bond acceptors, and it indicates their remarkable bonding pattern. Analyzing it through Figure1 and Table1, it signifies a strong interaction network between Baloxavir marboxil and hemagglutinin protein. Interacting with the receptor at 16, 10 and 11 sites orderly, the closest interaction distances between the drug and protein are orderly found to be within the range of 1.223-1.796, 0.97-1.54 and 1.376-1.899 with an average score of 1.393±0.138, 1.443±0.177 and 1.503±0.183 respectively. And, it clearly indicates a significantly closer interaction of Baloxavir marboxil in comparison to the other drugs. The drug, recently reaching the market in 2018 (Roche, 2018), targets the conserved substructures and shows the lowest binding energy and Ki score of −9.32Kj/mol and 146.33nM respectively. However, as it still misses out on several ADME properties, there is still a need to improve it further, and the present report provides reliable data to quickly pursue it further.

## Conclusion

The presented study performs the sequence and structural analysis of the HA sequence AWK91180.1 in comparison to its 150 closest orthologues. The 62.641KDa protein is estimated to be significantly stable and a set of 267 residues or 47.67% residues are found to be significantly conserved. Among the seven screened drugs, Baloxavir marboxil is found to be the most potent drug molecule with a binding affinity and Ki score of −9.32Kcal/mol and 146.33nM.

Excavating the interaction profiles of all the ligands, it is found that GLU407 and ILE276 residues interact with all the drugs, and are simply defining the interaction fingerprint for these molecules. Moreover, THR74, GLU76, ARG100 and PRO101 are found to have crucial interactions with the top two drug molecules: Baloxavir marboxil and Oleanane Acid, and are probably the key sites for a most potent drug molecule. The study builds a theoretical framework to build the most potent drug. To develop such a molecule, active against all diverse HA strains, the study is certainly the need of the hour.

## Notes

### Competing Interest Statement

The authors have declared no competing interest.

